# Metagenomic data reveal diverse fungal and algal communities associated with the lichen symbiosis

**DOI:** 10.1101/2020.03.04.966853

**Authors:** Hayden Smith, Francesco Dal Grande, Lucia Muggia, Rachel Keuler, Pradeep K. Divakar, Felix Grewe, Imke Schmitt, H. Thorsten Lumbsch, Steven D. Leavitt

## Abstract

Lichens have traditionally been considered the symbiotic phenotype from the interactions of a single fungal partner and one or few photosynthetic partners. However, the lichen symbiosis has been shown to be far more complex and may include a wide range of other interacting organisms, including non-photosynthetic bacteria, accessory fungi, and algae. In this study, we analyzed metagenomic shotgun sequences to better characterize lichen mycobiomes. Specifically, we inferred the range of fungi associated within lichen thalli from five groups of lichens – horsehair lichens (mycobiont=*Bryoria* spp.), shadow lichens (taxa in Physciaceae), rock posies (*Rhizoplaca* spp.), rock tripes (*Umbilicaria* spp.), and green rock shields (*Xanthoparmelia* spp.). Metagenomic reads from the multi-copy nuclear ribosomal internal transcribed spacer region, the standard DNA barcode region for fungi, were extracted, clustered, and used to infer taxonomic assignments. Our data revealed diverse lichen-associated mycobiomes, and closely related mycobionts tended to have more similar mycobiomes. Many of the members of the lichen-associated mycobiomes identified here have not previously been found in association with lichens. We found little evidence supporting the ubiquitous presence of Cystobasidiales yeasts in macrolichens, although reads representing this putative symbiotic partner were found in samples of horsehair lichens, albeit in low abundance. Our study further highlights the ecosystem-like features of lichens, with partners and interactions far from being completely understood. Future research is needed to more fully and accurately characterize lichen mycobiomes and how these fungi interact with the major lichen components – the photo- and mycobionts.

## 1. Introduction

Lichens have been iconic examples of symbiosis for the past 150 years (Honegger 2000). While the lichen was originally defined as a symbiotic relationship between a single fungus, the mycobiont, and a single or few species of green algae or cyanobacteria, the photobiont, studies have shown this is overly simplistic. It wasn’t until the late 20_th_ century that in vitro studies began to look at other fungi as potentially lichen-associated organisms rather than mere contaminants (Petrini et al. 1990, Crittenden et al. 1995, Girlanda et al. 1997).

Advances in sequencing technologies have allowed for a deeper investigation into the diversity of the lichen symbiosis, providing evidence that lichens are often composed of several fungal and green algal/cyanobacterial species forming a thallus with associated non-photosynthetic bacteria. Photobiont diversity can be shaped by reproductive and dispersal strategies of the mycobiont (Cao et al. 2015, Steinova et al. 2019), geography (Muggia et al. 2014, Werth and Sork 2014, Leavitt et al. 2015b), growth substrate (Bačkor et al. 2010, Leavitt et al. 2013b, Muggia et al. 2014) and macroclimate (Lu et al. 2018, Singh et al. 2018). The diversity of photobionts has been only recently explored by environmental DNA metabarcoding approaches and has focused on species within the Mediterranean basin to date (Moya et al. 2017, Dal Grande et al. 2018). In contrast to high-throughput sequencing approaches, traditional and largely applied DNA barcoding using Sanger sequencing was able to detect only the principal photobiont in the thalli (Paul et al. 2018). Additionally, many studies show that lichens are supported by a consortium of bacteria (Bates et al. 2011) that may change with substrate, altitude, and geography (Cardinale et al. 2012, Hodkinson et al. 2012, Fernandez-Brime et al. 2019). Potential functions of bacterial microbiomes include providing the host with nutrients, as well as protective and growth-regulating functions (Cernava et al. 2017). Furthermore, study have also shown carbon exchange between lichen green algae and non-photosynthetic bacteria (Kono et al. 2017).

The lichen mycobiome – the fungal communities superficially associated and within the lichen thallus – can be made up of symptomatic lichenicolous fungi (Lawrey and Diederich 2003) and endolichenic fungi (Arnold et al. 2009, U’Ren J et al. 2010, Muggia et al. 2016). Lichenicolous fungi growing on lichen thalli, may or may not be parasitic, and can influence their host’s morphology (Lawrey and Diederich 2003, U’Ren J et al. 2010, Fleischhacker et al. 2015). While some studies have found patterns in the lichen-associated mycobiome – for example, changing with altitude (Zhang et al. 2015, Wang et al. 2016) – others have found little specificity between the lichen host and its associated mycobiome (Fleischhacker et al. 2015, Fernandez-Mendoza et al. 2017, Yu et al. 2018).

Recently basidiomycete yeasts have been called into question as a potential symbiotic partner in the lichen symbiosis with the discovery of Cystobasidiomycetes (Basidiomycota, Pucciniomycotina) in the cortices of lichens (Spribille et al. 2016). The presence of this group of fungi was previously discovered in association with two genera in the lichen-forming family Parmeliaceae, *Hypogymnia* and *Usnea* by (Millanes et al. 2016), who clarified the phylogenetic position and the monophyly of two lichen-inhabiting species which were accommodated in the new genus *Cyphobasidium*. Later (Černajová and Škaloud 2019) found Cystobasidiomycete yeasts in 95% of *Cladonia* specimens collected across Europe, though they were suggested to be either part of a superficial biofilm or living within the thallus without associating with the cortex itself. In contrast, Lendemer et al. (2019) found them in just nine of the 339 species investigated. There remains a question of how abundant and specific cystobasidiomycetes are in lichen assemblages, as well as how consistent the mycobiome might be among different lichen-forming fungal species.

In terms of lichen photobionts, intrathalline photobiont diversity, e.g. multiple photobionts species within a single lichen thallus, has previously been observed in a number of lichen symbioses (Muggia et al. 2013, Dal Grande et al. 2014, Moya et al. 2017, Škaloud et al. 2018). In some cases, algae with different physiological performances are ever‐present in lichen thalli potentially facilitating the success of these lichens in a wide range of habitats and geographic areas and/or in changing environmental conditions. However, Sanger sequencing has been shown to consistently fail to effectively generate DNA sequence data from lichen specimens when multiple *Trebouxia* lineages occur within a single lichen thallus (Paul et al. 2018), potentially biasing the perspective of lichen photobiont associations. The prevalence of intrathalline photobiont diversity in lichens remains unclear, impacting our understanding of its ecological and evolutionary significance.

As lichens are a model of symbiosis, there is a need to better characterize their microbial partners and associations. Therefore, we used existing datasets of metagenomic shotgun sequences in an attempt to: (1) characterize the lichen mycobiomes across multiple, phylogenetically distinct lichen groups, (2) assess the prevalence of basidiomycete yeast, a putative symbiotic partner in some lichen symbioses, and (3) investigate the potential for multiple species-level *Trebouxia* algal lineages within a single lichen thallus.

## 2. Materials and Methods

### 2.1 Taxon sampling

Our sampling focused on five morphologically distinct lichens groups – (i) rock posy lichens – the *Rhizoplaca melanophthalma* group (Fig. 1A & B), (ii) green rock shield lichens – *Xanthoparmelia* spp. (Fig. 1C & D), (iii) rock tripe lichens – *Umbilicaria* spp., (iv) horse hair lichens – *Bryoria* spp., and (iv) representatives from the mycobiont family Physciaceae (shadow lichens) (Table 1; Fig. 1). Rock posy lichens were represented by three distinct forms, all occurring in western North America: the vagrant taxon *Rhizoplaca arbuscula* (Fig. 1B; n=3), the vagrant/erratic taxon *R. melanophthalma* subsp. *crispa* (n=3), and the rock-dwelling taxon *R. melanophthalma* (Fig. 1A; n=3) (Leavitt et al. 2013a). Green rock shield lichens were also represented by three distinct forms occurring in western North America: vagrant forms representing *Xanthoparmelia* aff. *chlorochroa* (Fig. 1D; n=3), isidiate (vegetative reproductive propagules) forms (Fig 1C; n=3), and the sexually reproducing taxon *X. neocumberlandia* (n=3) (Leavitt et al. 2011). Rock tripe lichens were represented by two species collected in Spain, *U. hispanica* (3 populations) and *U. pustulata* (Fig 1G; 2 populations). For the rock tripe lichens, each sample represents metagenomic reads from a pooled population – 100 lichen thalli/population – (Dal Grande et al. 2017), rather than reads from an individual lichen thallus. Horsehair lichens were represented by two species, *Bryoria fremontii* (Fig. 1H; n=3) and *B. tortuosa* (n=3) (Velmala et al. 2009). The fungal family Physciaceae was represented by *Mobergia calculiformis* (*Leavitt 16-697* [BRY-C]), *Physcia* sp. (*Leavitt 17-611* [BRY-C]), *Physciella* sp. (*Leavitt 17586* [BRY-C]), *Oxnerella* sp. (*Leavitt 17-611* [BRY-C]), and *Rinodina* sp. (*Leavitt 16-665* [BRY-C]). For rock posy lichens, green rock shield lichens, and representatives of Physciaceae, specimens were collected in dry conditions, with subsamples for molecular study removed within 24 h of collection and frozen at −20 °C until DNA extraction. Sampling of horse hair and rock trips lichens were reported previously in (Spribille et al. 2016b) and (Dal Grande et al. 2017, Dal Grande et al. 2018), respectively.

**Figure 1.**
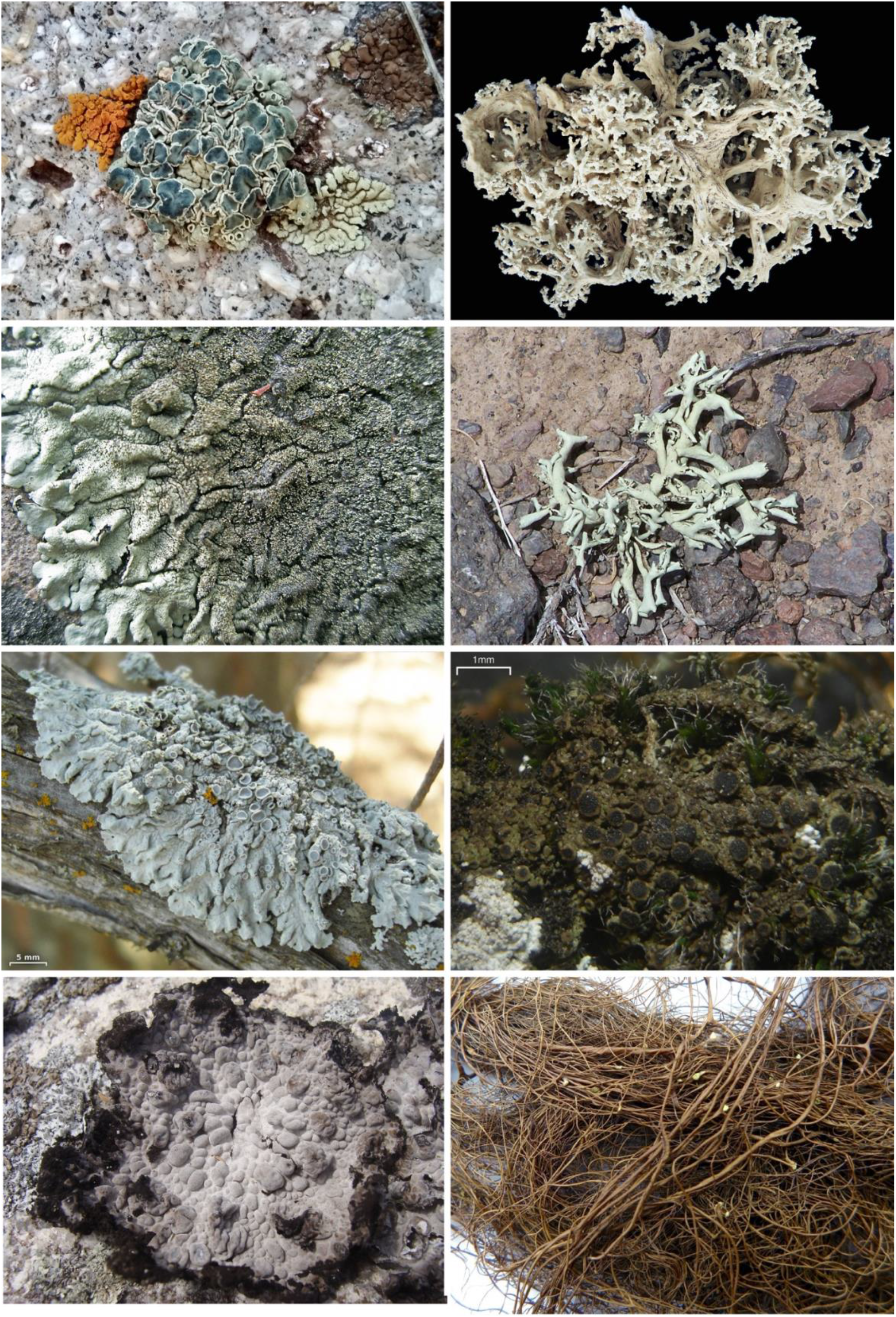
Examples of lichens groups considered in this study, including rock posies (**A** & **B**), green rock shields (**C** & **D**), shadow lichens (**E** & **F**), rock tripe (**G**), and horsehair lichens (**H**). **A**, *Rhizoplaca melanophthalma* – field image from La Sal Mountain Range, Utah, USA. **B**, *Rhizoplaca arbuscula* – collected from Lemhi Valley, Idaho, USA, voucher *Leavitt 18-1017* (BRY-C). **C**, *Xanthoparmelia* cf. *mexicana* – field image from Snake Range, Nevada, USA. **D**, *Xanthoparmelia* aff. *chlorochroa* – field image from Awapa Plateau, Utah, USA. **E**, *Physcia biziana* – field image from vicinity of Santa Fe, New Mexico (Hollinger 2492). **F**, *Rinodina* sp. (I need to track this down). **G**, *Umbilicaria pustulata* (I need to track this down). **H**, *Bryoria fremontii* (I need to track this down). Note: the name listed for each lichen of the mycobiont (main fungal partner) and does not account for the range of potential other associated symbionts.

#### Metagenomic sequencing

Metagenomic short reads used in this study originated from a range of sources and sequencing methods (Table 1). Metagenomic reads from rock posy lichens (*Rhizoplaca* spp.) were initially reported in (Leavitt et al. 2016, Leavitt et al. 2019) and are available in NCBI’s Short Read Archive under project PRJNA576709. For newly generated metagenomic reads from green rock shield lichens (*Xanthoparmelia* spp.) and representatives of Physciaceae, total genomic DNA was extracted from a small portion of lichen thalli (comprised of the mycobiont, photobiont, and other associated microbes) using the E.Z.N.A. Plant DNA DS Mini Kit (Omega Bio-Tek, Inc., Norcross, GA, USA) following the manufacturers’ recommendations. Total genomic DNA was prepared following the standard Illumina whole genome sequencing (WGS) library preparation process using Adaptive Focused Acoustics for shearing (Covaris), followed by an AMPure cleanup step. The DNA was then processed with the NEBNext Ultra™ II End Repair/dA-Tailing Module end-repair and the NEBNext Ultra™ II Ligation Module (New England Biolabs) while using standard Illumina index primers. Libraries were pooled and sequenced with the HiSeq 2500 sequencer in high output mode at the DNA Sequencing Center, Brigham Young University, Provo, Utah, USA, using either 250 cycle paired-end reads or 300 cycle paired-end reads. Reads from green rock shield lichens (*Xanthoparmelia* spp.) and representatives from the mycobiont family Physciaceae are available in NCBI’s Short Read Archive under project PENDING. The reads from the horsehair lichens are distinct in that these are transcriptomic reads (Spribille et al. 2016b), and we aimed to extract by-catch reads representing the internal transcribed spacer region (ITS). For the rock tripe lichens, each sample represents metagenomic reads from a pooled population (Pool-seq) – 100 lichen thalli/population – (Dal Grande et al. 2017), rather than reads from an individual lichen thallus.

#### Sequence Analysis

All reads were filtered using TRIMMOMATIC v0.33 (Bolger et al. 2014) before mapping to remove low quality reads and/or included contamination from Illumina adaptors using the following parameters: ILLUMINACLIP; LEADING:3; TRAILING:3; SLIDINGWINDOW:4:15; and MINLEN:36.

Previous studies have used assembled metagenomic contigs (Keepers et al. 2019) or mapping fungal reads to a fungal protein database (LaBonte et al. 2018) to provide crucial insight into fungal diversity in lichens and deciduous trees. Given the expected low coverage for fungi potentially co-occurring with a lichen thallus in short reads generated for this study, we chose to focus on the well-known repeat region which includes the standard fungal DNA barcode, the internal transcribed spacer (ITS) region of the nuclear ribosomal DNA (nrDNA) (Schoch et al. 2012). Across fungi, nrDNA copy number has been shown to vary considerably, ranging from tens to over 1400 copies per genome (Lofgren et al. 2019, Bradshaw et al. 2020). Furthermore, a comparatively robust and well-curated ITS database exists for fungi (Nilsson et al. 2019).

For reference ITS sequences, we used the UNITE QIIME v.8 dynamic release for fungi (Nilsson et al. 2019), filtered to include only sequences between 300 to 800 base pairs (reduced from 70,512 to 69,872 ITS sequences). Following recommendations by QIIME 2 developers, flanking regions, e.g., portions of the18S and/or 28S, with ITS sequences in the UNITE database were retained to reduce erroneous classifications when using the naïve Bayes classifier (https://doi.org/10.7287/peerj.preprints.27295v2). The UNITE ITS database was supplemented with all Cystobasidiomycetes ITS sequences reported in (Spribille et al. 2016). All sampled lichens are reported to associate with members of the genus *Trebouxia* as the primary lichen photobiont. In addition to assessing fungal diversity in short reads generated from lichen thalli, we also included representative sequences for each of the *Trebouxia* OTUs circumscribed in (Leavitt et al. 2015). Although lichens are known to associate with a broader range of algae than the core photobionts (Muggia et al. 2013), we did not assess accessory algae outside of *Trebouxia*.

For each metagenomic library, reads were mapped back to the composite ITS database using the Geneious read mapper in Geneious Prime (Kearse et al. 2012), implementing ‘Medium-Low Sensitivity / Fast’ sensitivity, iterated two times and saving all successfully mapped reads. Exploratory analyses with other read mapping approaches consistently recovered lower quantities of successfully mapped reads (data not shown). For each sample, metagenomic reads successfully mapped to the ITS references were imported into QIIME 2 (Bolyen et al. 2019). Reads were dereplicated using Vsearch ‘dereplicate-sequences’ (Rognes et al. 2016), implementing default settings. The dereplicated sequences were clustered into de novo OTUs at a 97% similarity in Vsearch using ‘cluster-features-de-novo’ (McDonald et al. 2012, Rognes et al. 2016). A naïve Bayes taxonomic classifier was trained using the same ITS reference library in QIIME 2 (Bokulich et al. 2018). The OTUs were then taxonomically classified using the trained naive Bayes trainer using QIIME 2 ‘feature-classifier classify-sklearn’ at a 0.95 confidence level to minimize false positives, with all other settings at default (McKinney 2010, Pedregosa et al. 2011, Bokulich et al. 2018).

Of the estimated 2.2 to 3.8 million fungal species, only 3–8% are currently named (Hawksworth and Lücking 2017), and a much smaller portion are represented in available DNA reference libraries. Exploratory analyses of our lichen mycobiome data revealed poor taxonomic resolution below class levels for the majority of OTUs inferred here. Therefore, fungal OTUs that were classified at the class level were retained and others with less taxonomic resolution were excluded. Classification of fungal OTUs generated from reads mapped to the reference ITS database was summarized using the QIIME ‘Taxa Barplot’ feature (Caporaso et al. 2010). Data were managed, analyzed and visualized in R (R Core Team, 2019) using ggplot2 (Wickham 2016) and tidyr (Wickham et al. 2019). To assess the similarity of lichen mycobiomes within and among phylogenetically distinct mycobionts, a principle component analysis (PCA) was performed on the class-level taxonomic classification using tidyr (Wickham et al. 2019), with the command ‘prcomp’. While formal species-level taxonomy in the lichen photobiont *Trebouxia* remains woefully inadequate (Muggia et al. *in review*), DNA sequence data representing a wide range of putatively species-level lineages, with accompanying provisional names, is available (Leavitt et al. 2015). For *Trebouxia* (photobiont) OTUs, the classified reads were filtered at the ‘species’ level, based on the 69 putative species-level OTUs from Leavitt et al. (2015), using QIIME ‘taxa filter-table’ command to determine the range of *Trebouxia* diversity occurring within each sample. All code used in this experiment is provided as supplementary file S1.

## 3. Results

Between 0.41 and 3.68% of metagenomic reads were mapped back to the ITS reference library (Table 1). The primary lichen symbionts, the mycobiont and photobiont, accounted for ca. 50% of all ITS reads extracted from the metagenomic data on average (Fig. 2A). The relative abundance of ITS reads representing the mycobiont (inferred at the class level, e.g., Lecanoromycetes) was between 5.20% to 80.31% of ITS reads, with an average relative abundance of 40%. The relative abundance of reads from the photobiont, *Trebouxia* spp., comprised between 0.68% to 35.09% of ITS reads, with an average relative abundance of ca. 10%.

**Figure 2.**
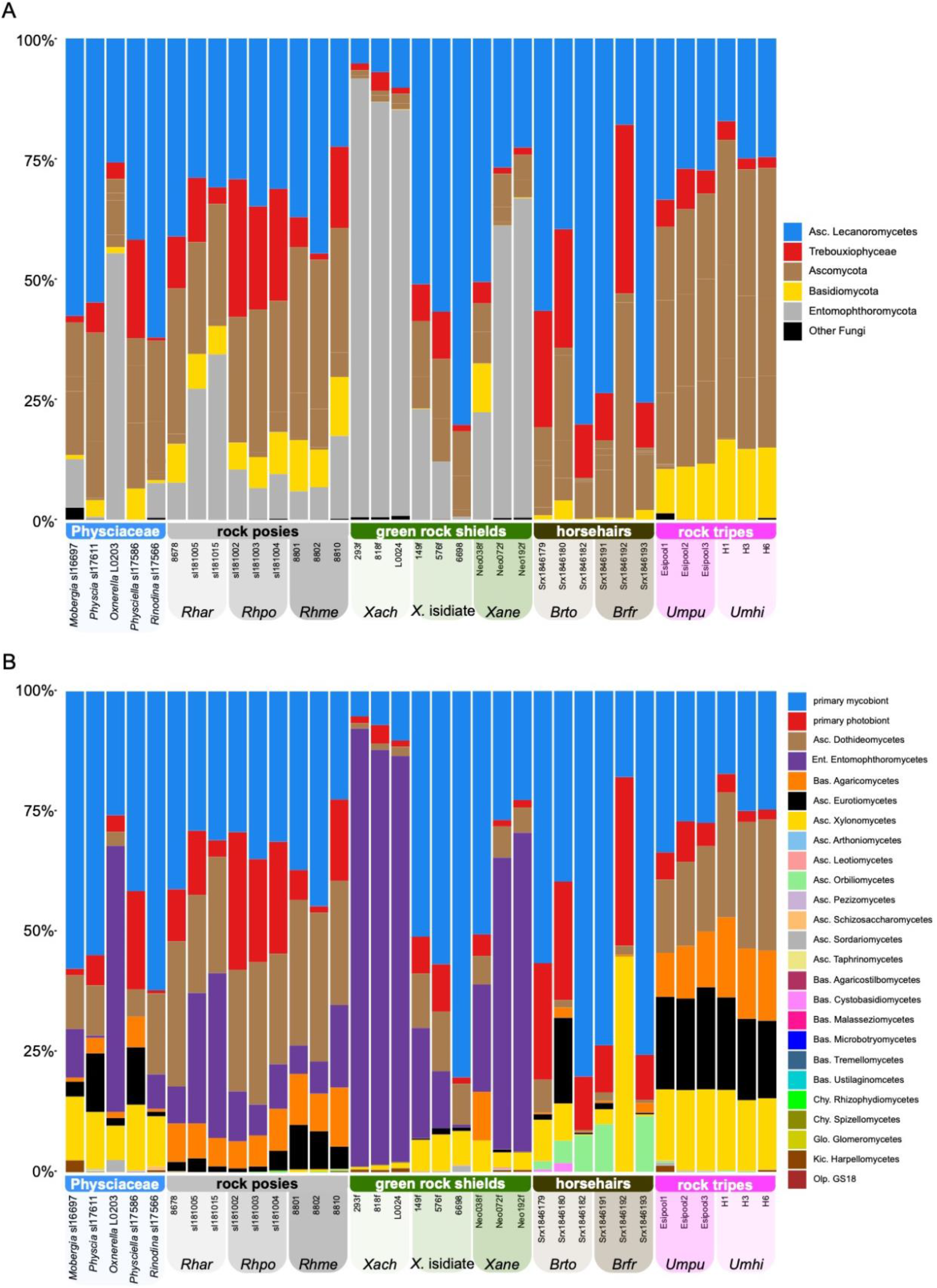
Overview of lichen symbionts and associated fungi inferred from data from the internal transcribed spacer region extracted from metagenomic shotgun sequencing short reads sequenced from lichen thalli representing five different groups of lichens. **A**, proportion of reads assigned to the lichen symbionts – mycobiont (shown in blue) and photobiont (in red) – and other major fungal lineages. **B**, same as in panel ‘A’, but major fungal lineages are broken down and shown at the class level. The five samples at the far left represent lichens associating with different members of the mycobiont family Physciaceae – ‘shadow lichens’; ‘Rock posies’ – in grey – are represented by three species in the mycobiont genus *Rhizoplaca*, with the first three specimens representing *R. arbuscula* (‘*Rhar*’), the following three samples are *R. melanophthalma* subsp. *crispa* (vagrant forms – ‘*Rhpo*’), and the final three samples represent *R. melanophthlama* (rock-dwelling, fertile forms – ‘*Rhme*’); ‘green rock shields’ – in green – are represented by three groups in the mycobiont genus *Xanthoparmelia*, with the first three samples representing *X.* aff. *chlorochroa* (asexual, vagrant forms – ‘*Xach*’), the next three samples represent isidiate, rock-dwelling forms (‘*X.* isidiate’), and the final three samples represent *X. neocumberlandia* (fertile, rock-dwelling forms – ‘*Xane*’); ‘horsehairs’ – in brown – are represented by two species in the mycobiont genus *Bryoria*, with the first three samples represent *B. tortuosa* (‘*Brto*’) and the last three, *B. fremontii* (‘*Brfr*’); ‘rock tripes’ – in pink – are represented by two groups in the mycobiont genus *Umbilicaria*, with the first three specimens representing *U. pustulata* (‘*Uspu*’)and the last three, *U. hispanica* (‘*Ushi*’). See supplementary file 1 for a full list of sampled lichens.

Lichen-associated fungi made up a large fraction of metagenomic reads, representing a total of 22 different fungal classes (Fig. 2B). Both in terms of abundance and diversity, Ascomycota OTUs were most frequently recovered and represented by 10 classes, excluding the mycobiont class Lecanoromycetes, followed by Basidiomycota represented by seven classes. Chytridiomycota (represented by two classes), Glomeromycota (one class), and Kickxellomycota (one class) were found in low abundance and diversity (at the class level) (abbreviated Chy., Glo., and Kic., respectively, in Fig. 3). Overall, reads from Cystobasidiomycete yeasts were poorly represented in extracted ITS reads, found in only 5 of the 35 samples. Notably, ITS by-catch from the *Bryoria fremontii* transcriptomic data from which lichen-associated yeasts were first reported in the cortex, resulted in the highest abundance of reads potentially representing cystobasidiomycete yeasts, with an average relative abundance of 0.7% of the ITS reads in the three *B. fremontii* samples. In the remaining two samples with evidence of Cystobasidiomycete yeasts, *Physcia biziana* and one sample of *Xanthoparmelia chlorochroa* (818F), had an average relative abundance of 0.03%.

**Figure 3.**
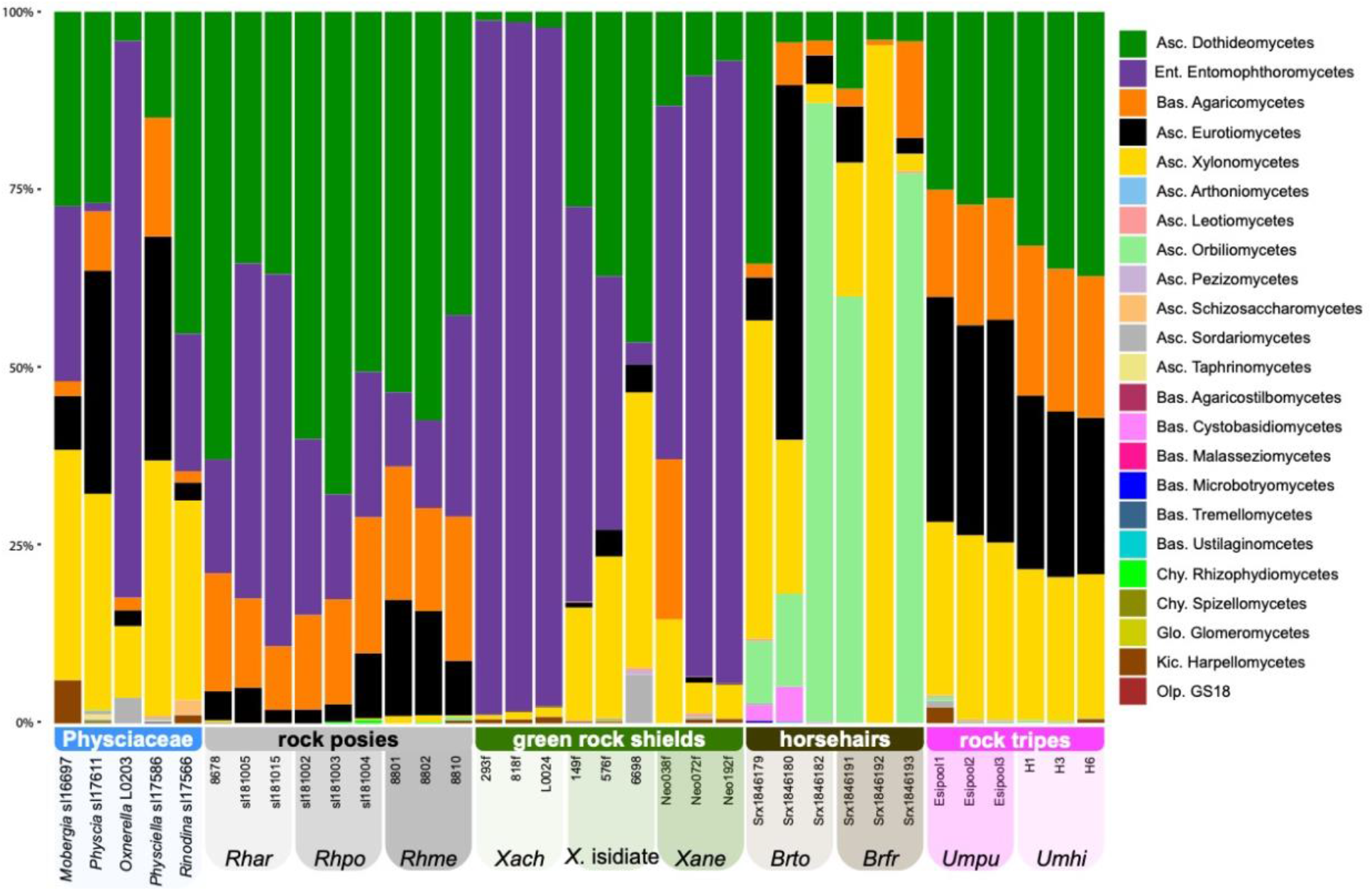
Inferred membership of lichen mycobiomes (at class level) inferred from data from the internal transcribed spacer region extracted from metagenomic shotgun sequencing short reads sequenced from lichen thalli (main lichen symbionts are excluded). The five samples at the far left represent lichens associating with different members of the mycobiont family Physciaceae – ‘shadow lichens’; ‘rock posies’ – in grey – are represented by three species in the mycobiont genus *Rhizoplaca*, with the first three specimens representing *R. arbuscula* (‘*Rhar*’), the following three samples are *R. melanophthalma* subsp. *crispa* (vagrant forms – ‘*Rhpo*’), and the final three samples represent *R. melanophthlama* (rock-dwelling, fertile forms – ‘*Rhme*’); ‘green rock shields’ – in green – are represented by three groups in the mycobiont genus *Xanthoparmelia*, with the first three samples representing *X.* aff. *chlorochroa* (asexual, vagrant forms – ‘*Xach*’), the next three samples represent isidiate, rock-dwelling forms (‘*X.* isidiate’), and the final three samples represent *X. neocumberlandia* (fertile, rock-dwelling forms – ‘*Xane*’); ‘horsehairs’ – in brown – are represented by two species in the mycobiont genus *Bryoria*, with the first three samples represent *B. tortuosa* (‘*Brto*’) and the last three, *B. fremontii* (‘*Brfr*’); ‘rock tripes’ – in pink – are represented by two groups in the mycobiont genus *Umbilicaria*, with the first three specimens representing *U. pustulata* (‘*Uspu*’)and the last three, *U. hispanica* (‘*Ushi*’). See supplementary file 1 for a full list of sampled lichens.

Closely related mycobionts tended to have more similar mycobiomes (Fig. 4). The PCA revealed a general pattern of mycobiome similarity among samples representing mycobiont species, and relatively high levels of similarity among mycobiont congeners (Figs. 3 & 4). Differences in lichen mycobiomes are most distinct among different genera of lichen-forming fungi.

**Figure 4.**
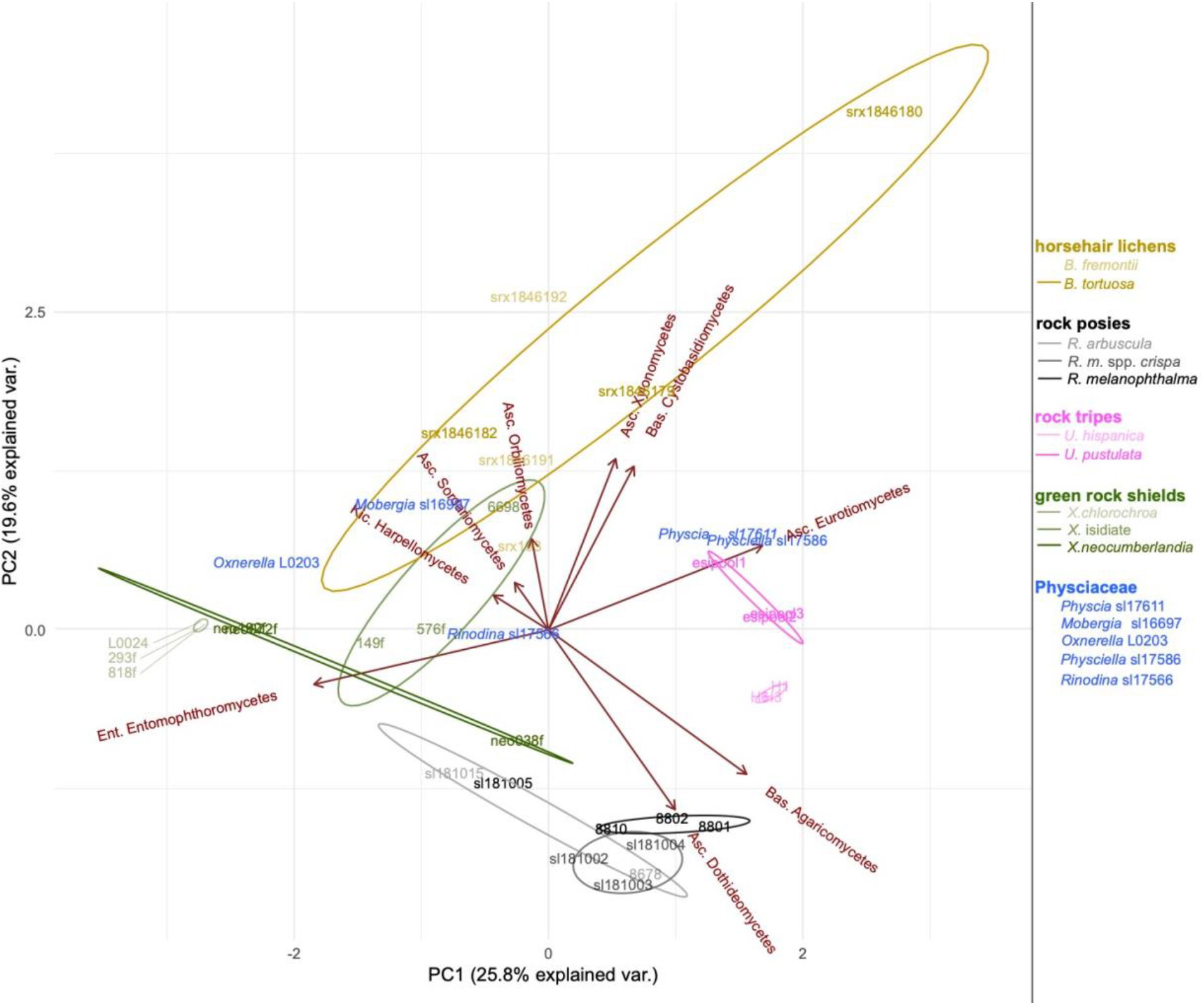
Principal component analysis of lichen mycobiome diversity.

Evidence supporting intrathalline *Trebouxia* photobiont diversity was observed in 16 of the 29 samples (*Umbilicaria* samples not considered – see methods above) (Fig. 5). Thalli from representatives of Physciaceae and *Rhizoplaca* (rock posy lichens) consistently contained a dominant *Trebouxia* lineage with >90% relative abundance. Green rock shield lichens (*Xanthoparmelia* spp.) associated with a wider range of *Trebouxia* species, with evidence of multiple *Trebouxia* species occurring within an individual lichen thallus. Two of the six *Bryoria* samples also provided evidence of multiple *Trebouxia* species occurring within individual thalli. Intrathalline photobiont diversity in *Umbilicaria pustulata* and U. *hispanica* is described in detail in (Paul et al. 2018). Here we report photobiont diversities within populations of *Umbilicaria* spp. (each sample represents 100 pooled individual thalli from a single population).

**Figure 5.**
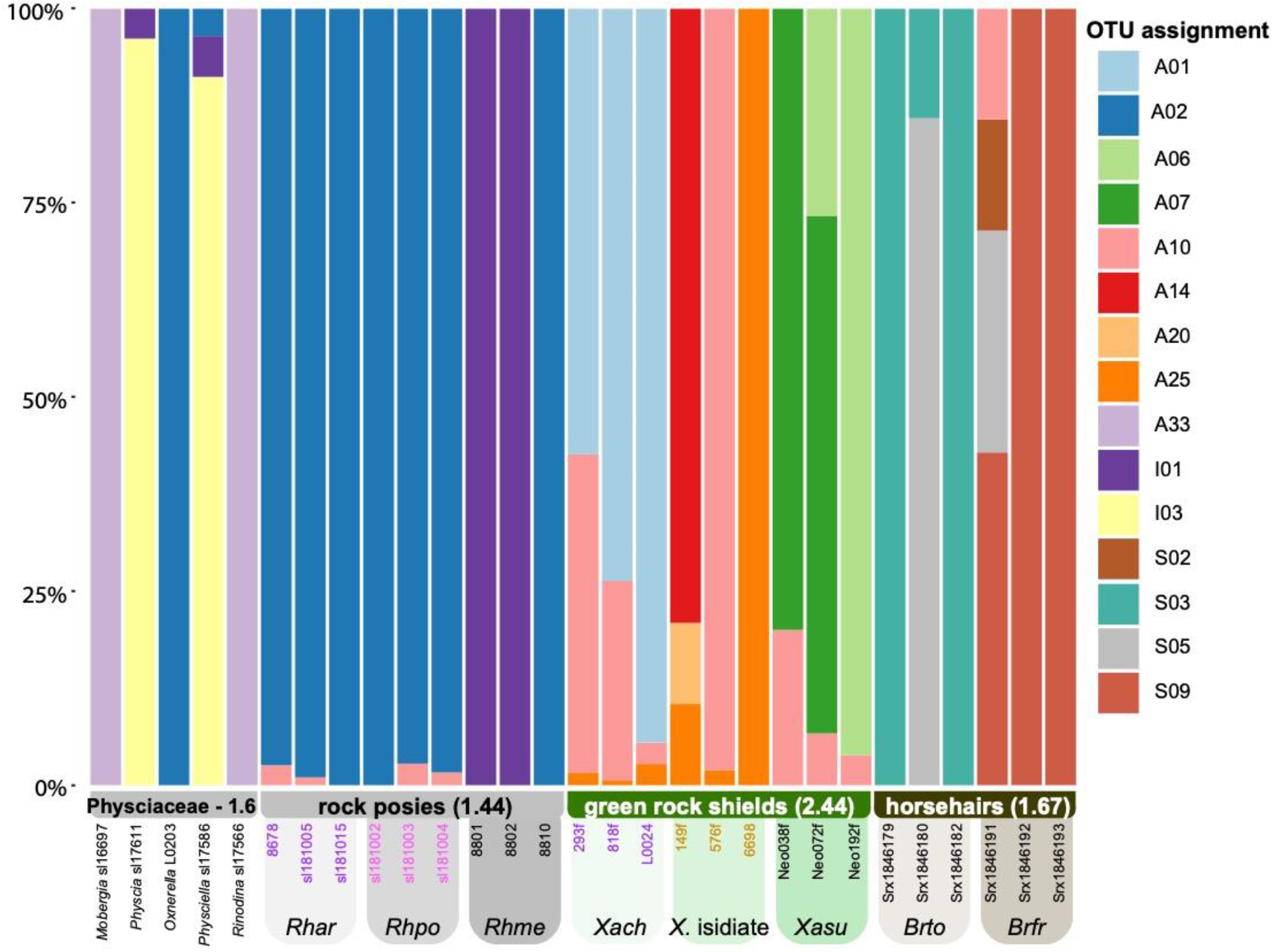
Assessment of intrathalline *Trebouxia* (photobiont) diversity in four lichen groups. *Trebouxia* OTUs nomenclature follows Leavitt et al. 2015. ‘Rock posies’ – in grey – are represented by three species in the mycobiont genus *Rhizoplaca*, with the first three specimens representing *R. arbuscula* (‘*Rhar*’), the following three samples are *R. melanophthalma* subsp. *crispa* (vagrant forms – ‘*Rhpo*’), and the final three samples represent *R. melanophthlama* (rock-dwelling, fertile forms – ‘*Rhme*’); ‘green rock shields’ – in green – are represented by three groups in the mycobiont genus *Xanthoparmelia*, with the first three samples representing *X.* aff. *chlorochroa* (asexual, vagrant forms – ‘*Xach*’), the next three samples represent isidiate, rock-dwelling forms (‘*X.* isidiate’), and the final three samples represent *X. neocumberlandia* (fertile, rock-dwelling forms – ‘*Xane*’); and ‘horsehairs’ – in brown – are represented by two species in the mycobiont genus *Bryoria*, with the first three samples represent *B. tortuosa* (‘*Brto*’) and the last three, *B. fremontii* (‘*Brfr*’).

## 4. Discussion

The broad range of organisms involved in lichen symbioses has recently been highlighted, including diverse algae (Muggia et al. 2013, Moya et al. 2017), non-photosynthetic bacteria (Cardinale et al. 2006, Grube et al. 2009, Hodkinson and Lutzoni 2009), and broad range of fungal lineages (Lawrey and Diederich 2003, Spribille et al. 2016, Tuovinen et al. 2019). Using data mining of fungal ITS reads from metagenomic shotgun sequences of lichen thalli, we provide a coarse snapshot of unexpectedly diverse lichen-associated mycobiomes (Fig. 3). The accessory fungi accounted for a significant proportion of ITS reads extracted from metagenomic shotgun sequencing data (Fig. 2B), spanning multiple phyla – dominated by Ascomycota and Basidiomycota but with representatives from Entomophthoromycota, Chytridiomycota, Glomeromycota, and Kickxellomycota. While a number of the class-level lineages inferred from metagenomic ITS reads are known to associate with lichens, e.g., Agaricomycetes, Dothideomycetes, Eurotiomycetes, and Sordariomycetes, other classes do not include fungi previously known to associate with lichens, e.g. Entomophthoromytes. In contrast to recent studies highlighting the role of two basidiomycete lineages in some lichen symbioses, *Tremella* (Tuovinen et al. 2019) and Cystobasidiomycete yeasts (Spribille et al. 2016), these were recovered only sporadically and in very low abundance in our samples. Nonetheless, these basidiomycete fungi have often been reported as lichenicolous, growing on a number of lichen hosts (Diederich 1996, Millanes et al. 2016). Below we discuss the potential implications of our findings and potential ways to move forward.

The relative importance of host versus environment in determining the diversity of the lichen mycobiome is poorly understood. However, lichen mycobiomes appear to comprise stable and transient guilds, which to some extent correlate with the ecological conditions of the lichen habitats. (Fernández-Mendoza et al. 2017) proposed three ecological components of lichen mycobiomes: (i) generalist taxa common to the environmental pool of bio- and saprotrophic fungi, (ii) lichenicolous and endolichenic fungi specific to each genus/species, and (iii) species which disperse and possibly germinate on, among, and within lichen thalli, but do not play a definite ecological role in the lichen community. Our results indicate that closely related mycobionts tend to have more similar mycobiomes (Fig. 4), even in cases where distinct lichens commonly co-occur, e.g. green rock shields (mycobiont = *Xanthoparmelia* spp.) and rock posies (mycobiont = *Rhizoplaca* spp.). These data support the perspective that a significant component of the lichenicolous and endolichenic fungal community are specific to different mycobiont genera/species.

Broadly speaking, Sordariomycetes and Leotiomycetes are frequently recovered from lichens occurring in humid, temperate, boreal environments, and Antarctic environments, representing lineages closely related to plant endophytes (Arnold et al. 2009, U’Ren J et al. 2010, Yu et al. 2018). In contrast, Dothideomycetes and Eurotiomycetes are more frequently associated with rock-inhabiting lichens (Muggia and Grube 2018). In rock-inhabiting lichens, the lichen-associated fungi are usually melanized fungi comprising unknown and known hyphomycetous lineages which show close affinities to some symptomatic lichenicolous fungi, extremotolerant rock-inhabiting fungi from oligotrophic environments and to plant and animal pathogenic black yeasts (Muggia et al. 2016, Muggia and Grube 2018). These fungi are widely known as black fungi because they accumulate melanins in their cell walls, which enable them to grow in oligotrophic environments and resist multiple abiotic stresses, such as high doses of radiation, desiccation, temperature extremes (Gostinčar et al. 2009). Black fungi are, therefore, usually recognized as (poly)extremotolerant organisms.

In our study, different lichen genera tended to associate with distinct fungal communities (Fig. 3). These relationships appear to be consistent across relatively broad geographic areas, at least for some lichens. Our results indicated that the mycobiomes of green rock shield (mycobiont = *Xanthoparmelia* spp.) and rock posy (mycobiont = *Rhizoplaca* spp.) populations occurring across western North America were strikingly different (Fig. 3). While disparate morphologies of rock posy lichens had relatively consistent mycobiomes, even in specimens collected across geographically distinct populations, differences in mycobiome communities of green rock shield lichens with different morphologies and reproductive strategies were observed (Fig. 3). However, within green rock shield lichens, vagrant (obligately unattached specimens), rock-dwelling isidiate (reproducing via specialized asexual propagules), and rock-dwelling sexually reproducing forms tended to associate with distinct fungal communities, albeit with limited sample sizes. Additional research will be required to more fully assess if distinct mycobiomes, or core subsets of the mycobiome, within lichen groups are maintained across geographic and ecological distances. If differing core mycobiome communities are found in association with distinct mycobionts, at what level does this specificity exist, e.g., mycobiont species, genera, etc.? Directed experimental design and broader sampling will be required to determine how lichen mycobiomes are structured at different evolutionary scales relative to the predominant mycobiont.

When investigating the potential for photobiont (*Trebouxia* spp.) diversity within a single lichen thallus, our results suggest that a single lichen thallus of some lichen groups, e.g., rock posies (mycobiont = *Rhizoplaca* spp.) and shadow lichens (members of the mycobiont family Physciaceae), tend to associate with a single/one dominant *Trebouxia* lineage. For rock tripe lichens (mycobiont = *Umbilicaria* spp.), (Paul et al. 2018) observed a single pattern of a single dominant *Trebouxia* lineage per thallus. However, the metagenomic reads from rock tripe lichens used in the present study were generated from multiple lichen thalli pooled into a single population per site, and we were unable to corroborate these results reported. In contrast, it appears that green rock shield lichen (mycobiont = *Xanthoparmelia*) thalli consistently harbor multiple, distinct *Trebouxia* lineages. A previous study characterizing *Trebouxia* diversity associating with members of the mycobiont family Parmeliaceae also demonstrated distinct patterns of photobiont association between *Rhizoplaca* spp. and *Xanthoparmelia* spp., with *Xanthoparmelia* spp. associating with a much wider range of photobionts than *Rhizoplaca* spp. (Leavitt et al. 2015). Furthermore, the two mycobiont genera consistently associated with distinct *Trebouxia* lineages with very little overlap, and these results were corroborated by our findings (Fig. 5). By explicitly taking the potential for intrathalline photobiont diversity into consideration, we anticipate novel insight into different strategies of lichen symbiosis.

While our study provides novel insight into lichen symbioses and impetus for future research, there are a number of methodological limitations that potentially bias the results presented here. Metagenomic reads from lichen-forming fungi are expected to be dominated by reads from the major lichen symbionts, the myco- and photobionts (Pizarro 2019), and other eukaryotic microbial diversity associated with lichen thalli is likely found in much lower abundance in metagenomic short read data. Therefore, here we opted to target fungal reads from the multi-copy nuclear ribosomal cistron (nrDNA) in order to identify fungi that might be found in low relative abundance and likely overlooked using single copy regions and metagenomic binning approaches. Furthermore, with portions of the nrDNA are highly conserved across fungi, we focused on the highly variable internal transcribed spacer region (ITS) due to the high variability and well-curated reference database (Schoch et al. 2012, Nilsson et al. 2019). However, nrDNA copy number varies by orders of magnitude across fungi, from tens to over 1400 copies per genome (Lofgren et al. 2019, Bradshaw et al. 2020). Therefore, the relative abundance of fungal groups inferred in this study (e.g., Fig. 3) does not accurately depict true relative abundance of lichen-associated fungi given the potential for a very wide range of nrDNA copy number of these fungi.

Another source of potential bias is from the bioinformatic pipeline implemented here. Even using relatively well-established pipelines of ITS amplicon-based metagenomic reads, bioinformatics analysis pipelines have been shown to vary greatly in their relative performance and accuracy in characterizing fungi from metagenomic data (Anslan et al. 2018). We would anticipate that the data mining approach implemented in this study may have introduced a number of unexpected and difficult to identify artifacts, ranging from potentially over- and underrepresenting different fungal lineages to erroneous taxonomic assignments. The impact of these potential methodological limitations is not clear. For example, in the present study, a significant proportion of reads from green rock shield lichens were assigned to the class Entomophthoromytes, a lineage that has not previously been found in association with lichens. Whether the inferred prevalence of Entomophthoromytes is biased by copy number variation of the nrDNA, an artifact of read mapping to the UNITE database, etc., or accurately represents a novel finding is unclear. Furthermore, only a small fraction of the estimate 2.2–3.8 million fungal species are represented in currently available curated databases. Therefore, in fungal metabarcoding studies, a large proportion of operational taxonomic units (OTUs) cannot be identified to any meaningful taxonomic, and these unclassifiable species hypotheses, or ‘dark taxa’, remain problematic in metagenomic studies of fungi (Nilsson et al. 2019).

Taken together, our results highlight, on the one hand, the presence of a highly diverse, seemingly lichen host-specific mycobiome, and on the other hand, the risk of applying overly simplistic techniques – such as phylum rank classifications – to tackle the diversity of these lichen-associated fungal communities.

## Acknowledgements

We gratefully acknowledge the fruitful discussion with Toby Spribille, Diane Haughland, and Trevor Goward. This research was supported by College of Life Sciences at Brigham Young University, Provo, Utah, USA. We thank Ed Wilcox, DNA Sequencing Center, Brigham Young University, Provo, Utah, USA, for help with sequencing.

## References

Anslan S, Nilsson RH, Wurzbacher C, Baldrian, P, Leho, T, Bahram, M (2018) Great differences in performance and outcome of high-throughput sequencing data analysis platforms for fungal metabarcoding. MycoKeys, pp 29–40.

Arnold AE, Miadlikowska J, Higgins KL, Sarvate SD, Gugger P, Way A, Hofstetter V, Kauff F, Lutzoni F (2009) A phylogenetic estimation of trophic transition networks for ascomycetous fungi: are lichens cradles of symbiotrophic fungal diversification? Syst Biol 58, 283–297.

Bačkor M, Peksa O, Škaloud P, Bačkorová M (2010) Photobiont diversity in lichens from metal-rich substrata based on ITS rDNA sequences. Ecotoxicology and Environmental Safety 73, 603–612.

Bates, ST, Cropsey GW, Caporaso JG, Knight R, Fierer N (2011) Bacterial communities associated with the lichen symbiosis. Appl Environ Microbiol 77, 1309–1314.

Bokulich NA, Kaehler, BD, Rideout, JR, Dillon, M, Bolyen, E, Knight, R, Huttley, GA, Gregory Caporaso J (2018) Optimizing taxonomic classification of marker-gene amplicon sequences with QIIME 2’s q2-feature-classifier plugin. Microbiome 6, 90.

Bolger, AM, Lohse M, Usadel B (2014) Trimmomatic: a flexible trimmer for Illumina sequence data. Bioinformatics 30, 2114–2120.

Bolyen E, Rideout JR, Dillon MR, et al. (2019) Reproducible, interactive, scalable and extensible microbiome data science using QIIME 2. Nat Biotechnol 37, 852–857.

Bradshaw M, Grewe F, Thomas A, Harrison CH, Lindgren H, Muggia L, St Clair LL, Lumbsch HT, Leavitt SD (2020) Characterizing the ribosomal tandem repeat and its utility as a DNA barcode in lichen-forming fungi. BMC Evolutionary Biology 20, 2.

Cao S, Zhang F, Liu C, Hao Z, Tian Y, Zhu L, Zhou Q (2015) Distribution patterns of haplotypes for symbionts from *Umbilicaria esculenta* and *U. muehlenbergii* reflect the importance of reproductive strategy in shaping population genetic structure. BMC Microbiol 15, 212.

Caporaso JG, Kuczynski J, Stombaugh J et al. (2010) QIIME allows analysis of high-throughput community sequencing data. Nat Methods 7, 335–336.

Cardinale M, Grube M, Castro Jr JV, Muller H, Berg G (2012) Bacterial taxa associated with the lung lichen *Lobaria pulmonaria* are differentially shaped by geography and habitat. FEMS Microbiol Lett 329, 111–115.

Cardinale M, Puglia AM, Grube M (2006) Molecular analysis of lichen-associated bacterial communities. FEMS Microbiology Ecology 57, 484–495.

Černajová I, Škaloud P (2019) The first survey of Cystobasidiomycete yeasts in the lichen genus *Cladonia*; with the description of *Lichenozyma pisutiana* gen nov, sp nov. Fungal Biology 123, 625–637.

Cernava T, Erlacher A, Aschenbrenner IA, Krug L, Lassek C, Riedel K, Grube M, Berg G (2017) Deciphering functional diversification within the lichen microbiota by meta-omics. Microbiome 5, 82.

Crittenden PD, David JC, Hawksworth DL, Campbell FS (1995) Attempted isolation and success in the culturing of a broad spectrum of lichen-forming and lichenicolous fungi. The New Phytologist 130, 267–297.

Dal Grande F, Alors D, Divakar PK, Bálint M, Crespo A, Schmitt I (2014) Insights into intrathalline genetic diversity of the cosmopolitan lichen symbiotic green alga *Trebouxia decolorans* Ahmadjian using microsatellite markers. Molecular Phylogenetics and Evolution 72, 54–60.

Dal Grande F, Rolshausen G, Divakar PK, Crespo A, Otte J, Schleuning M, Schmitt I (2018) Environment and host identity structure communities of green algal symbionts in lichens. New Phytologist 217, 277–289.

Dal Grande F, Sharma R, Meiser A, Rolshausen G, Büdel B, Mishra B, Thines M, Otte J, Pfenninger M, Schmitt I (2017) Adaptive differentiation coincides with local bioclimatic conditions along an elevational cline in populations of a lichen-forming fungus. BMC Evolutionary Biology 17, 93.

Diederich P, 1996 The lichenicolous heterobasidiomycetes. Bibliotheca Lichenologica 61, 1–198.

Fernandez-Brime S, Muggia L, Maier S, Grube M, Wedin M (2019) Bacterial communities in an optional lichen symbiosis are determined by substrate, not algal photobionts. FEMS Microbiol Ecol 95.

Fernandez-Mendoza F, Fleischhacker A, Kopun T, Grube M, Muggia L (2017) ITS1 metabarcoding highlights low specificity of lichen mycobiomes at a local scale. Mol Ecol 26, 4811–4830.

Fernández-Mendoza F, Fleischhacker A, Kopun T, Grube M, Muggia L (2017) ITS1 metabarcoding highlights low specificity of lichen mycobiomes at a local scale. Molecular Ecology 26, 4811–4830.

Fleischhacker A, Grube M, Kopun T, Hafellner J, Muggia L (2015) Community analyses uncover high diversity of lichenicolous fungi in alpine habitats. Microb Ecol 70, 348–360.

Girlanda M, Isocrono D, Bianco C, Luppi-Mosca A (1997) Two foliose lichens as microfungal ecological niches. Mycologia 89, 531–536.

Gostinčar C, Grube M, De Hoog S, Zalar P, Gunde-Cimerman N (2009) Extremotolerance in fungi: evolution on the edge. FEMS Microbiology Ecology 71, 2–11.

Grube M, Cardinale M, de Castro Jr JV, Muller H, Berg G (2009) Species-specific structural and functional diversity of bacterial communities in lichen symbioses. ISME J 3, 1105–1115.

Hawksworth DL, Lücking R (2017) fungal diversity revisited: 22 to 38 million species. Microbiol Spectr 5.

Hodkinson B, Lutzoni F (2009) A microbiotic survey of lichen-associated bacteria reveals a new lineage from the Rhizobiales. Symbiosis 49, 163–180.

Hodkinson BP, Gottel NR, Schadt CW, Lutzoni F (2012) Photoautotrophic symbiont and geography are major factors affecting highly structured and diverse bacterial communities in the lichen microbiome. Environ Microbiol 14, 147–161.

Honegger R, 2000 Simon Schwendener (1829-1919) and the dual hypothesis of lichens. The Bryologist 103, 307–313.

Kearse M, Moir R, Wilson A, Stones-Havas S, Cheung M, Sturrock S, Buxton S, Cooper A, Markowitz S, Duran C, Thierer T, Ashton B, Meintjes P, Drummond A (2012) Geneious Basic: An integrated and extendable desktop software platform for the organization and analysis of sequence data. Bioinformatics 28, 1647–1649.

Keepers KG, Pogoda CS, White KH, Anderson Stewart CR, Hoffman JR, Ruiz AM, McCain,CM, Lendemer JC, Kane NC, Tripp EA (2019) Whole genome shotgun sequencing detects greater lichen fungal diversity than amplicon-based methods in environmental samples. Frontiers in Ecology and Evolution 7.

Kono M, Tanabe H, Ohmura Y, Satta Y, Terai Y (2017) Physical contact and carbon transfer between a lichen-forming *Trebouxia* alga and a novel *Alphaproteobacterium*. Microbiology 163, 678–691.

LaBonte NR, Jacobs J, Ebrahimi A, Lawson S, Woeste K (2018) Data mining for discovery of endophytic and epiphytic fungal diversity in short-read genomic data from deciduous trees. Fungal Ecology 35, 1–9.

Lawrey JD, Diederich P (2003) Lichenicolous fungi: interactions, evolution, and biodiversity. The Bryologist 106, 80–120.

Leavitt SD, Fernández-Mendoza F, Pérez-Ortega S, Sohrabi M, Divakar PK, Vondrák J, Lumbsch HT, St. Clair LL (2013)a Local representation of global diversity in a cosmopolitan lichen-forming fungal species complex (*Rhizoplaca*, Ascomycota). Journal of Biogeography 40, 1792–1806.

Leavitt SD, Nelsen MP, Lumbsch HT, Johnson LA, St Clair LL (2013b) Symbiont flexibility in subalpine rock shield lichen communities in the Southwestern USA. The Bryologist 116, 149–161.

Leavitt SD, Grewe F, Widhelm T, Muggia L, Wray B, Lumbsch HT (2016) Resolving evolutionary relationships in lichen-forming fungi using diverse phylogenomic datasets and analytical approaches Scientific Reports 6, 22262.

Leavitt SD, Johnson LA, Goward T, St Clair LL (2011) Species delimitation in taxonomically difficult lichen-forming fungi: An example from morphologically and chemically diverse *Xanthoparmelia* (Parmeliaceae) in North America. Molecular Phylogenetics and Evolution 60, 317–332.

Leavitt SD, Keuler R, Newberry CC, Rosentreter R, St Clair LL (2019) Shotgun sequencing decades-old lichen specimens to resolve phylogenomic placement of type material. Plant and Fungal Systematics 64, 237–247

Leavitt SD, Kraichak E, Nelsen MP, Altermann S, Divakar PK, Alors D, Esslinger TL, Crespo A, Lumbsch HT (2015) Fungal specificity and selectivity for algae play a major role in determining lichen partnerships across diverse ecogeographic regions in the lichen-forming family Parmeliaceae (Ascomycota). Mol Ecol 24, 3779–3797.

Lendemer JC, Keepers KG, Tripp EA, Pogoda CS, McCain CM, Kane NC (2019) A taxonomically broad metagenomic survey of 339 species spanning 57 families suggests cystobasidiomycete yeasts are not ubiquitous across all lichens. Am J Bot 106, 1090–1095.

Lofgren LA, Uehling JK, Branco S, Bruns TD, Martin F, Kennedy PG (2019) Genome-based estimates of fungal rDNA copy number variation across phylogenetic scales and ecological lifestyles. Mol Ecol 28, 721–730.

Lu J, Magain N, Miadlikowska J, Coyle JR, Truong C, Lutzoni F (2018) Bioclimatic factors at an intrabiome scale are more limiting than cyanobiont availability for the lichen-forming genus *Peltigera*. Am J Bot 105, 1198–1211.

McDonald D, Clemente JC, Kuczynski J, Rideout JR, Stombaugh J, Wendel D, Wilke A, Huse S, Hufnagle J, Meyer F, Knight R, Caporaso JG (2012) The biological observation matrix (BIOM) format or: how I learned to stop worrying and love the ome-ome. Gigascience 1, 7.

McKinney W (2010) Data Structures for Statistical Computing in Python.

Millanes AM, Diederich P, Wedin M (2016) *Cyphobasidium* gen nov, a new lichen-inhabiting lineage in the Cystobasidiomycetes (Pucciniomycotina, Basidiomycota, Fungi). Fungal Biology 120, 1468–1477.

Moya P, Molins A, Martínez-Alberola F, Muggia L, Barreno E (2017) Unexpected associated microalgal diversity in the lichen *Ramalina farinacea* is uncovered by pyrosequencing analyses. PLOS ONE 12, e0175091.

Muggia L, Fleischhacker A, Kopun T, Grube M (2016) Extremotolerant fungi from alpine rock lichens and their phylogenetic relationships. Fungal Diversity 76, 119–142.

Muggia L, Grube M (2018) Fungal diversity in lichens: from extremotolerance to interactions with algae. Life 8, 15.

Muggia L, Pérez-Ortega S, Kopun T, Zellnig G, Grube M (2014) Photobiont selectivity leads to ecological tolerance and evolutionary divergence in a polymorphic complex of lichenized fungi. Annals of Botany 114, 463–475.

Muggia L, Vancurova L, Škaloud P, Peksa O, Wedin M, Grube M (2013) The symbiotic playground of lichen thalli – a highly flexible photobiont association in rock-inhabiting lichens. FEMS Microbiology Ecology 85, 313–323.

Nilsson RH, Larsson K-H, Taylor AFS, Bengtsson-Palme J, Jeppesen TS, Schigel D, Kennedy P, Picard K, Glöckner FO, Tedersoo L, Saar I, Kõljalg U, Abarenkov K (2019) The UNITE database for molecular identification of fungi: handling dark taxa and parallel taxonomic classifications. Nucleic Acids Research 47, D259–D264.

Paul F, Otte J, Schmitt I, Dal Grande F (2018) Comparing Sanger sequencing and high-throughput metabarcoding for inferring photobiont diversity in lichens. Scientific Reports 8, 8624.

Pedregosa F, Varoquaux G, Gramfort A, Michel V, Thirion B, Grisel O, Blondel M, Prettenhofer P, Weiss R, Dubourg V, Vanderplas J, Passos A, Cournapeau D, Brucher M, Perrot M, Duchesnay É (2011) Scikit-learn: Machine learning in Python. Journal of Machine Learning Research 12, 2825–2830.

Petrini O, Hake U, Dreyfuss MM (1990) An analysis of fungal communities isolated from fruticose lichens. Mycologia 82, 444–451.

Pizarro D (2019) Metagenomic sequencing with new bioinformatics approaches to understand the evolution of lichen forming fungi (dissertation). Complutense University of Madrid Complutense University of Madrid, Madrid.

Rognes T, Flouri T, Nichols B, Quince C, Mahe F (2016) VSEARCH: a versatile open source tool for metagenomics. PeerJ 4, e2584.

Schoch CL, Seifert KA, Huhndorf S et al (2012) Nuclear ribosomal internal transcribed spacer (ITS) region as a universal DNA barcode marker for fungi. Proceedings of the National Academy of Sciences 109, 6241–6246. doi: 101073/pnas1117018109.

Singh G, Dal Grande F, Schnitzler J, Pfenninger M, Schmitt I (2018) Different diversification histories in tropical and temperate lineages in the ascomycete subfamily Protoparmelioideae (Parmeliaceae). Mycokeys, 1–19.

Škaloud P, Moya P, Molins A, Peksa O, Santos-Guerra A, Barreno E (2018) Untangling the hidden intrathalline microalgal diversity in *Parmotrema pseudotinctorum*: *Trebouxia crespoana* sp nov. The Lichenologist 50, 357–369.

Spribille T, Tuovinen V, Resl P, Vanderpool D, Wolinski H, Aime MC, Schneider K, Stabentheiner E, Toome-Heller M, Thor G, Mayrhofer H, Johannesson H, McCutcheon JP (2016) Basidiomycete yeasts in the cortex of ascomycete macrolichens. Science 353, 488–492.

Steinova J, Skaloud P, Yahr R, Bestova H, Muggi, L (2019) Reproductive and dispersal strategies shape the diversity of mycobiont-photobiont association in *Cladonia* lichens. Mol Phylogenet Evol 134, 226–237.

Tuovinen V, Ekman S, Thor G, Vanderpool D, Spribille T, Johannesson H (2019) Two basidiomycete fungi in the cortex of wolf lichens. Current Biology 29, 476–483e475.

U’Ren JM, Lutzoni F, Miadlikowska J, Arnold AE (2010) Community analysis reveals close affinities between endophytic and endolichenic fungi in mosses and lichens. Microb Ecol 60, 340–353.

Velmala S, Myllys L, Halonen P, Goward T, Ahti T (2009) Molecular data show that *Bryoria fremontii* and *B tortuosa* (Parmeliaceae) are conspecific. The Lichenologist 41, 231–242.

Wang Y, Zheng Y, Wang X, Wei X, Wei J (2016) Lichen-associated fungal community in *Hypogymnia hypotrypa* (Parmeliaceae, Ascomycota) affected by geographic distribution and altitude. Front Microbiol 7, 1231.

Werth S, Sork VL (2014) Ecological specialization in *Trebouxia* (Trebouxiophyceae) photobionts of *Ramalina menziesii* (Ramalinaceae) across six range-covering ecoregions of western North America. Am J Bot 101, 1127–1140.

Wickham H (2016) ggplot2: Elegant Graphics for Data Analysis Springer-Verlag New York

Wickham H, Averick M, Bryan J, Chang W, McGowan L, François R, Grolemund G, Hayes A, Henry L, Hester J (2019) Welcome to the Tidyverse. Journal of Open Source Software 4, 1686.

Yu NH, Park SY, Kim JA, Park CH, Jeong MH, Oh SO, Hong SG, Talavera M, Divakar PK, Hur JS (2018) Endophytic and endolichenic fungal diversity in maritime Antarctica based on cultured material and their evolutionary position among Dikarya. Fungal Systematics and Evolution 2, 263–272.

Zhang T, Wei XL, Zhang YQ, Liu HY, Yu LY (2015) Diversity and distribution of lichen-associated fungi in the Ny-Alesund Region (Svalbard, High Arctic) as revealed by 454 pyrosequencing. Sci Rep 5, 14850.

